# Experimental monkeypox virus infection in rats

**DOI:** 10.1101/2024.12.04.626788

**Authors:** Barry Rockx, Lucien van Keulen, Wim H.M. van der Poel, Kevin K. Ariën, Koen Vercauteren, Sandra van de Water, José Harders-Westerveen, Paul J. Wichgers Schreur, Joke van der Giessen, Miriam Maas, Norbert Stockhofe, Laurens Liesenborghs

## Abstract

The global spread of Monkeypox virus (MPXV) clade IIb in 2022/2023 raised concerns about spillback into new animal reservoirs. Experimental inoculation of rats with MPXV resulted in skin lesions and viral shedding in the respiratory tract and skin. These findings suggest a potential role for rats in MPXV transmission.

---

Mpox is a zoonotic disease caused by monkeypox virus (MPXV) and is currently the most prevalent orthopoxviral infection in humans after the eradication of smallpox. In endemic areas (African region), MPXV circulates among a number of mammal species, including shrews, rodents and primates, although the exact reservoir remains unknown(1). Until a few years ago, outbreak were mainly caused by spillover events from the animal reservoir and sustained human-to-human transmission was relatively rare.

In July 2022, however, the World Health Organization (WHO) declared mpox a Public Health Emergency of International Concern (PHEIC) due to the global spread of Clade IIb, a subclade of Clade II (previously known as the West African Clade)(2). More recently, a similar scenario unfolded in Central Africa, involving Clade I (formerly known as the Central African Clade). Initially, Clade I was predominantly transmitted zoonotically, but in 2023, a subclade called Clade Ib started to spread through human-to-human transmission across multiple countries(3). This led the WHO to declare a second PHEIC in August 2024.

MPXV can infect a wide range of mammal species, however no known animal reservoir currently exists outside of Africa. A major concern with the 2022/2023 outbreak was that MPXV may enter a new host species and establish a new animal reservoir outside of Africa through spillback from infected humans to their pets or farm animals. These infected animals in turn may then become reservoirs and transmit the virus back to humans. While a recent case of an MPXV infected individual transmitting MPXV to an animal (a dog)(4), was reported, this may have been due to DNA contamination from human mpox cases(5). In addition, the detection of MPXV in waste water adds to the potential risk of brown rats (*Rattus norvegicus*)(6), which often live in sewers, becoming a reservoir for this virus.

The main objective of this study was to assess the susceptibility of rats to MPXV and characterize the disease progression, viral shedding and transmission potential in an effort to understand whether rats could potentially act as (intermediate) host and ultimately might become reservoirs of MPXV following the introduction in countries outside of the endemic region.

## The Study

In order to determine whether rats are susceptible to infection with MPXV, we experimentally inoculated groups of 6 female Wistar IGS rats (age 6 weeks; Charles River), with 10^7^ per mL tissue culture dose 50% (TCID_50%_) of a clinical isolate of MPXV clade IIb, isolated from a Belgian patient in 2022, or phosphate-buffered saline (PBS) for mock-control animals. Animals were inoculated via intranasal instillation (50µL) as well as by scarification between the shoulderblades following shaving of the skin (∼5 µL bifurcated needle, provided by RIVM, Bilthoven, the Netherlands), in the same animal. Animals were monitored for the development of clinical disease daily. Regular physical examinations were performed and included daily body weights, rectal temperature, blood draw, and collection of skin, oral, ocular, rectal, and nasal swabs. At 2 and 10 days post inoculation (dpi), a group of 6 infected and 2 uninfected control animals were euthanized to study tissue distribution of virus and associated pathology. The animal experiment was conducted in accordance with European regulations (EU directive 2010/63/EU) and the Dutch Law on Animal Experiments (Wod, ID number BWBR0003081). Permissions were granted by the Dutch Central Authority for Scientific Procedures on Animals (Permit Number: AVD40100202216563). All experimental protocols were approved by the Animal Ethics Committees of Wageningen Research.

Following inoculation, no weight loss or fever were observed until the end of the experiment (Figure 1A and B). However, starting 4 dpi, skin lesions developed at the site of scarification (Figure 1C). These lesions were detectable up to 11 dpi. No other clinical signs were observed.

**Figure 1.**
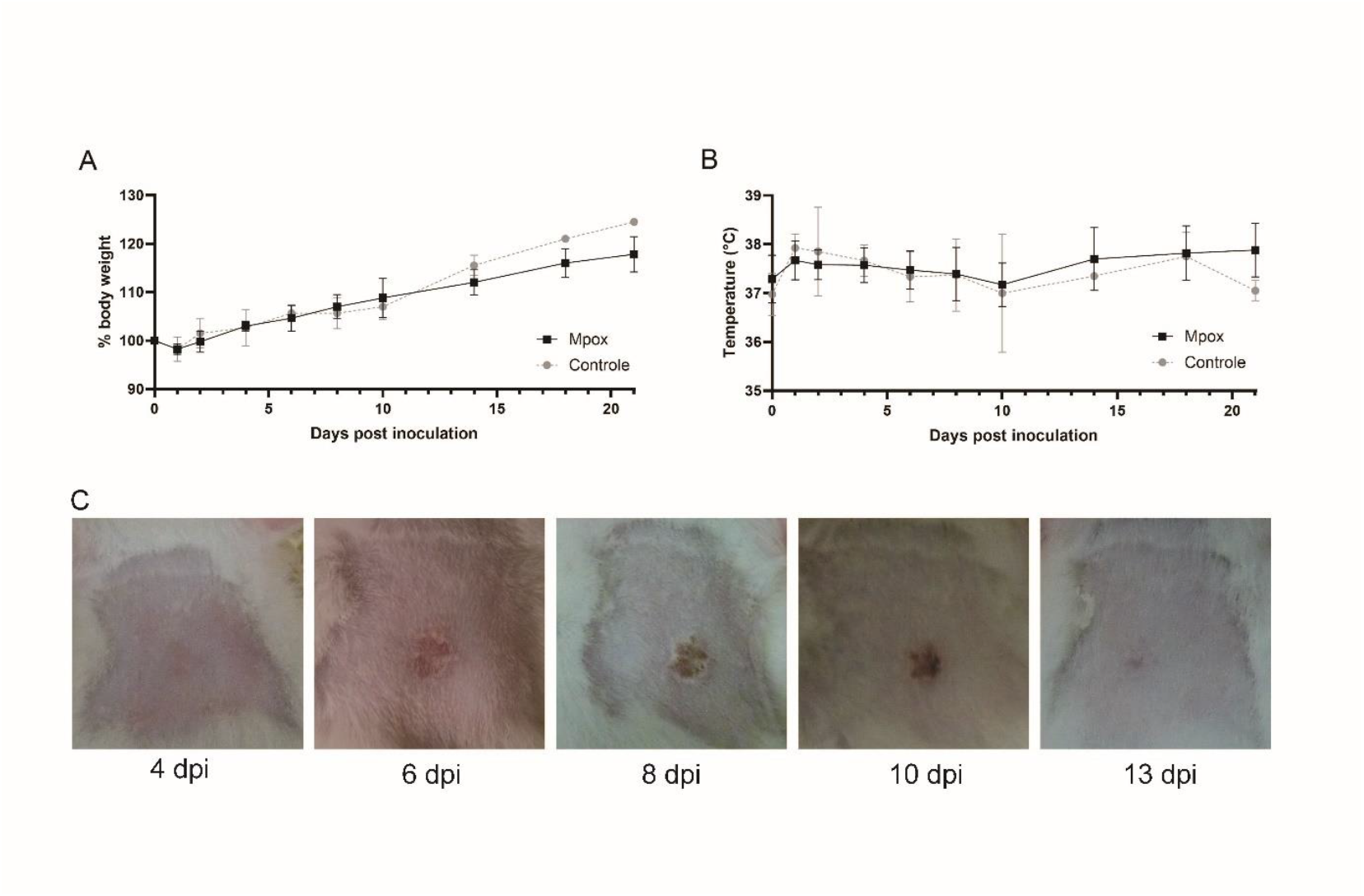
Clinical manifestations after MPXV inoculation in rats. Relative body weight (A) and rectal temperature (B) of rats inoculated with MPXV clade IIb (black) and mock controls (grey). The mean of each group is shown as solid line. Standard deviation shown as error bars. Photographic images of development of skin lesions on 4, 6, 8, 10 and 13 dpi from representative animal (C).

To determine whether infected animals shed virus, we examined the presence of viral genomic DNA in swabs of nose, throat, rectum and skin by qPCR(7). Low levels of viral genome were detected in nose swabs, in the majority of animals, on 6 dpi only (Figure 2A). Higher levels of viral genome were detected in throat swabs, with peak shedding on 6 or 8 dpi and no detectable genome by 10 dpi (Figure 2B). Infectious virus was only detected by virus isolation on Vero cells on 6 dpi, which corresponded with the highest genome loads. Although more variable, skin swabs were also positive for viral genome as early as 4 dpi (Figure 2C), and while the majority of animals was negative by 10 dpi, 1 animal remained positive until 15 dpi. Infectious virus was only detected in peak samples on 4 dpi. The absence of viral genome at earlier time points suggests that this was the result of active replication and not residual inoculum. Viral genome in rectal swabs was only detected in 1 animal and at a low relative load. All ocular swabs were tested negative (data not shown).

**Figure 2.**
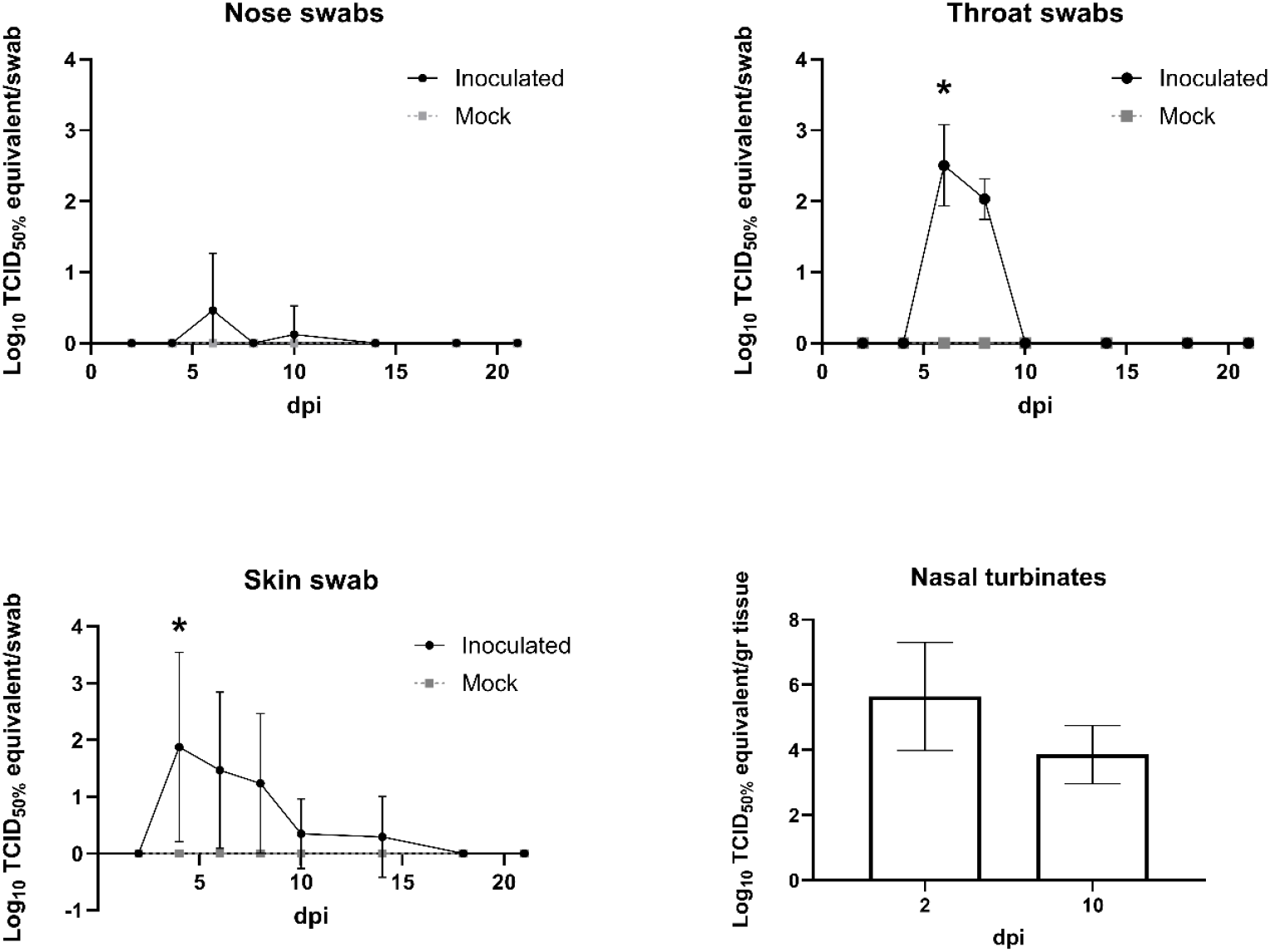
Kinetics of MPXV shedding and replication. Kinetics of shedding in nose (A), throat (B) and skin (C) swabs as determined by detection of viral genome. Viral replication is expressed as Log_10_ TCID_50%_ equivalent per swab. The mean viral load is shown as a solid black (inoculated) or grey (mock) line. * indicate isolation of infectious virus. Replication in nasal turbinates (D) as determined by detection of viral genome and expressed as Log_10_ relative viral load per gram tissue. The mean of each group is shown as a bar. Standard deviation is shown as error bars.

In addition to the detection of viral genome in swabs, nasal turbinates were positive in all animals sacrificed on 2 and 10 dpi (Figure 2D). While the lungs of 1 animal were positive 2 days dpi with a low viral genome load, the lungs of all other rats, as well as other tissues tested on 2 and 10dpi (blood, brain, skin, kidney, liver and spleen), remained negative.

Upon necropsy, enlarged cervical lymph nodes were observed in all infected animals at 10 dpi, but no other gross pathological findings were present at 2 or 10 dpi. Histopathological investigation showed that the cervical lymph nodes showed a reactive hyperplasia but no viral antigen could be detected immunohistochemically in the lymph nodes or nasal turbinates at 2 or 10 dpi. Skin lesions were not present at 2 dpi but were characterized by acanthosis, mononuclear inflammation and scattered necrotic keratinocytes in the epidermis at 10 dpi (Figure 3). Surprisingly, viral antigen was not detected in the skin at 2 or 10 dpi by immunohistochemistry (data not shown).

**Figure 3.**
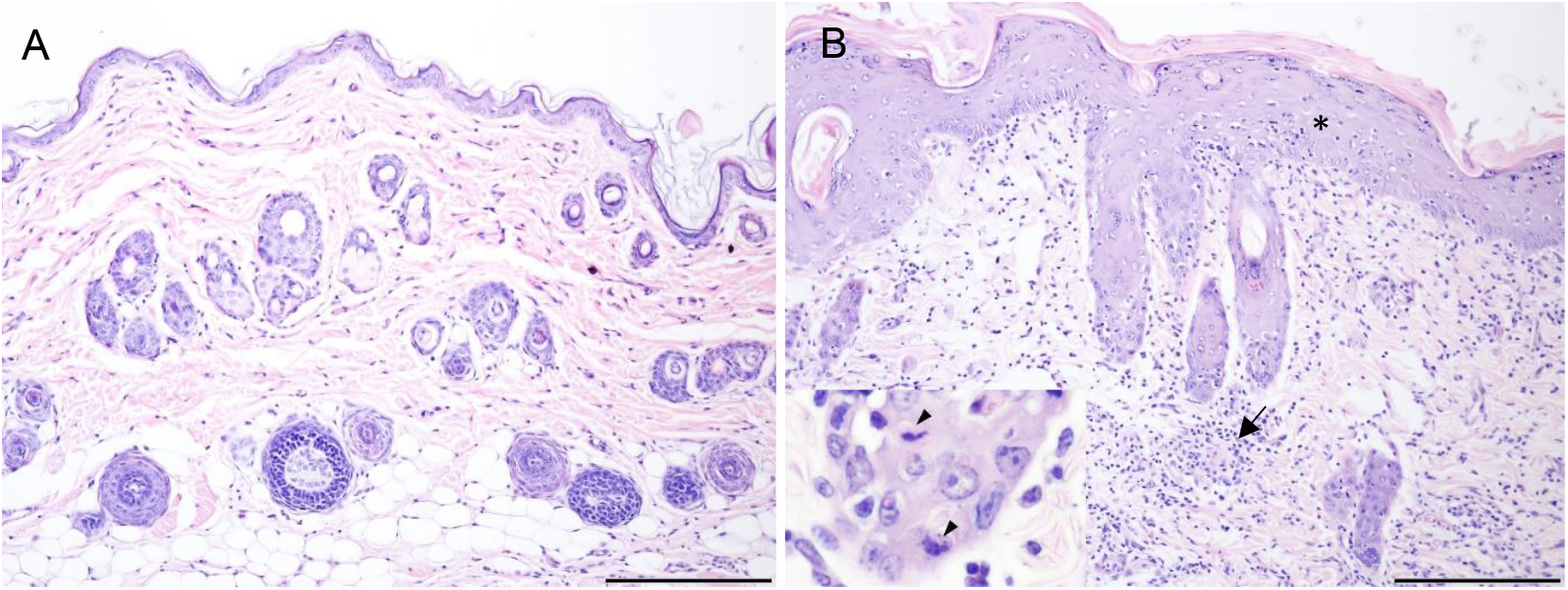
Histopathology of the rat skin, at the site of scarification (A) uninfected and (B) 10 days post infection with MPXV clade IIb. Notice the thickening of the epidermis (acanthosis, asterisk) and marked mononuclear inflammatory reaction in the dermis (arrow). Insert: higher magnification of the follicular epithelium showing necrotic keratinocytes (arrowheads). Haematoxylin and eosin staining, bar = 200 μm.

## Conclusions

The main objective of this study was to assess the susceptibility of rats to MPXV in an effort to determine whether rats could potentially act as (intermediate) hosts and ultimately might become reservoirs of MPXV following introduction in countries outside of the endemic region. A previous study had suggested that adult rats were not susceptible, while MPXV infection of newborn rats resulted in lethal disease within 6 dpi(8). Our study shows that 6 weeks old rats are susceptible to MPXV clade IIb infection. MPXV infection in rats results in an acute and self-limiting infection, characterized by a transient development of skin lesions. Infected rats shed infectious MPXV via the respiratory tract and skin lesions. One limitation is that these studies were performed with laboratory rats, rather than wild rats for biosafety purposes, and responses in outbred animal may differ. However, based on the acute self-limiting infection, lack of viremia, and lack of prolonged shedding as shown in this study, there is a low risk for rats to establish as a reservoir for MPXV.

In addition, in light of the ongoing multi-country outbreak of clade I MPXV(3), here we present a potential new infection and disease model that recapitulates the disease seen in the majority of human cases and can be used for preclinical evaluation of countermeasures against MPXV.

Dr. Rockx is a senior researcher at Wageningen Bioveterinary Research. His research interest focus on the pathogenesis and treatment of emerging and re-emerging zoonotic viruses in the context of One Health.

## Acknowledgement

We thank Koert Stittelaar at WUR, and Harry Vennema at RIVM for their input. We would like to thank our animal caretakers for their support during the animal experiment. This study was supported in part, by a Transnational Access (VetBioNet research services) grant from the ISIDORe infrastructure project (EU Horizon Europe, grant agreement number 101046133) and the Ministry of Health, Welfare and Sports in the Netherlands.

